# Systemic host inflammation induces stage-specific transcriptomic modification and slower maturation in malaria parasites

**DOI:** 10.1101/2021.08.18.456784

**Authors:** Lianne I.M. Lansink, Jessica A. Engel, Hyun Jae Lee, Megan S.F. Soon, Cameron G. Williams, Arya SheelaNair, Clara P.S. Pernold, Pawat Laohamonthonkul, Jasmin Akter, Thomas Stoll, Michelle M. Hill, Arthur M. Talman, Andrew Russell, Mara Lawniczak, Miles P. Davenport, David S. Khoury, Ashraful Haque

## Abstract

Maturation rate of malaria parasites within red blood cells (RBC) can be influenced by host nutrient status or circadian rhythm. Here, we observed in mice that systemic host inflammation, induced by lipopolysaccharide (LPS) conditioning or ongoing acute malaria infection, slowed the progression of a single cohort of parasites from one generation of RBC to the next. LPS-conditioning and acute infection both triggered substantial changes to the metabolomic composition of plasma in which parasites circulated. This altered plasma directly slowed parasite maturation in a manner that could not be rescued by supplementation, consistent with the presence of inhibitory factors. Single-cell transcriptomic assessment of mixed parasite populations, exposed to a short period of systemic host inflammation *in vivo*, revealed specific impairment in the transcriptional activity and translational capacity of trophozoites compared to rings or schizonts. Thus, we provide *in vivo* evidence of transcriptomic and phenotypic plasticity of asexual blood-stage *Plasmodium* parasites when exposed to systemic host inflammation.

## Introduction

Malaria, caused by infection with *Plasmodium* parasites, remains a significant global health problem, with ∼200 million cases and ∼400,000 deaths each year between 2015-2020^1^. *Plasmodium* parasites invade red blood cells (RBCs), and then mature, replicate, and burst out over ∼24, 48 or 72 hour periods, depending on the infecting species. Given each parasite produces up to 32 daughter merozoites, parasite populations can increase dramatically *in vivo*. Malaria symptoms occur during this asexual, cyclical stage, with disease severity and risk of death during *P. falciparum* infection correlating positively with parasite biomass^2^.

From a host perspective, slowing down parasite population growth constitutes an effective strategy for preventing or ameliorating severe disease. This can be achieved in multiple ways, for example by killing parasites with artemisinin-based antimalarial drugs^3,4^, or blocking their transit between RBCs with parasite-specific antibodies^5^. From a parasite perspective, there is theoretical benefit to adjusting the rate of progression through the asexual life-cycle, to avoid overwhelming a vulnerable host, at least until transmission is guaranteed.

*In vitro* experiments support the idea that certain molecules can control asexual stage biology in *Plasmodium* parasites. For example, isoleucine starvation and lactate supplementation have been reported to slow blood-stage *P. falciparum* maturation *in vitro*^6-8^. Lysophosphatidylcholine was reported to suppress cellular differentiation of asexual parasites into gametocytes^9^. While causal links remain to be provided *in vivo*, the prevailing model is that many different types of metabolite could directly influence asexual stage *Plasmodium* parasites. In recent years, *in vivo* evidence from our group and others supports the idea that *Plasmodium* parasites can sense and rapidly respond to dynamic changes within the mammalian host. Mancio-Silva *et al* demonstrated that *P. berghei* parasites upregulated expression of ∼180 genes within 6 hours of *in vivo* exposure to a calorie-restricted host – compared to control hosts fed *ad libitum*^10^. Rijo-Ferreira *et al* revealed altered periodicity of expression for ∼1000 *P. chabaudi* genes in hosts both harbouring a genetically altered circadian rhythm and being housed without light-dark cycling^11^. We showed the asexual life-cycle was plastic for *P. berghei* or *P. yoelii* parasites^12^, with the *P. berghei* parasite life-cycle extended from ∼24h to ∼40h by exposure to a pre-existing malaria infection^12^. Given this effect was lost in similarly infected immune-deficient *rag1*^*-/-*^ mice, we surmised that host-mediated responses, perhaps systemic inflammation had slowed maturation of parasites. Since parasitised RBCs (pRBC) circulate in plasma, we hypothesized in this study that alterations to plasma constituents directly mediate adjustments to parasite maturation rate.

In addition to changes in the host environment, we also aimed to understand the parasite changes contributing to delayed development in the context of host inflammation. Cellular change in eukaryotic cells, including protozoan parasites, can be detected by transcriptomic analysis. Bulk RNA-seq and microarray studies of synchronised *Plasmodium* parasites have yielded stage-specific gene expression signatures that have been crucial for mapping biological processes accompanying asexual parasite maturation^13,14^. More recently, single-cell RNA-seq (scRNA-seq) has enabled examination of *Plasmodium* parasites without requirement to synchronise or otherwise fractionate parasites into morphologically similar subsets^15^. These single-cell assessments have served to construct a “Malaria Cell Atlas” (MCA), which details transcriptomic changes as malaria parasites differentiate within mammalian and insect hosts^15^. In these reports, the asexual stages of the *P. berghei* MCA were generated *in vitro* in the absence of host immune pressure. Hence, the possible effect of host inflammation on individual asexual parasites remains unknown.

Here, we employed *P. berghei* ANKA infection or innate immune stimulation of mice in conjunction with unsupervised metabolomic assessment of plasma, *in vitro* culturing methods, and droplet-based scRNA-seq of parasites to reveal that systemic host inflammation triggers changes to the host plasma environment, which is rapidly sensed and responded to by parasites residing in RBC.

## Materials and Methods

### Mice

C57BL/6J mice were purchased from the Animal Resource Centre (Perth, Australia). C57BL/6J.*rag1*^*−/−*^ mice were bred at QIMR Berghofer Medical Research Institute. All mice were female between 6-12 weeks of age and were maintained under conventional conditions. This study was carried out in strict accordance with guidelines from The National Health and Medical Research Council of Australia. All animal procedures and protocols were approved (A02-633M and A1503-601M) and monitored by the QIMR Berghofer Medical Research Institute Animal Ethics Committee.

### *Plasmodium* infection

*Plasmodium berghei* ANKA parasites constitutively expressing high-levels of eGFP (for RBC adoptive transfer), or luciferase (for establishing acute infection, although bioluminescence was not utilised), were sourced and used as previously reported^12,16,17^. *Pb*A parasites were used after defrosting cryopreserved infected blood and a single *in vivo* passage in C57BL/6J mice. RBCs were collected from passage mice by cardiac puncture and used to infect with 10^5^ pRBCs via lateral tail vein injection.

### Adoptive transfer of fluorescently-labelled parasitised red blood cells

Adoptive transfer of fluorescently-labelled RBC was performed as previously described (^5,12^). Briefly, RBCs were collected from infected mice by cardiac puncture, washed twice in Ca^2+^- and Mg^2+^-free PBS, and stained with viable cell dye CellTrace™ Far Red (CTFR; Thermo Fisher), according to manufacturer’s instructions. CTFR-labelled RBCs were resuspended in 2 mL RPMI per donor mouse and injected in 200 μl volumes to recipient mice via lateral tail vein injection using 26G needles.

### TLR agonist treatment

Mice were treated with Saline (0.9%; Baxter), Lipopolysaccharide (LPS; 0.75 mg/mL) (Sigma-Aldrich), CpG 1826 (0.25 mg/mL) (Sigma-Aldrich) or Polyriboinosinic:polyribocytidylic acid (Poly I:C; 2 mg/mL) (InvivoGen) via intraperitoneal injection (200 μL per mouse) using 26G needles, two hours prior to adoptive transfer of CFTR-labelled RBCs.

### Flow cytometry

Flow cytometric analysis of pRBCs in peripheral blood was performed as previously described^12^. 1-2 drops (approx. 10-20 µL) of peripheral blood were collected by tail vein bleed and diluted in RPMI medium containing 5 U/mL heparin sulphate. Diluted blood was stained with Hoechst33342 (10 μg/mL) (Sigma-Aldrich) and Syto84 (5 μM) (Life Technologies) at room temperature in the dark for 30 minutes. Staining was quenched with 10x initial volume of ice-cold RPMI medium and samples acquired using a LSR II Fortessa analyzer (BD Biosciences) and analysed using FlowJo software version 10.7 (Treestar).

### Cytokine analysis

Serum or plasma cytokine levels were assessed using the BD CBA Mouse Inflammation kit (BD Biosciences), as per manufacturer’s instructions. Data were acquired on a BD LSR II Fortessa Analyzer (BD Biosciences) and analysed using FCAP Array Software version 3.0 (BD Biosciences).

### Plasma metabolomics

Two independent experiments were conducted, each with 6 mice per treatment group. Ice-cold butanol/methanol (1:1) containing 50 µg/mL antioxidant 2,6-di-tert-butyl-4-methylphenol (BHT) was added to each plasma sample, as well as a pooled quality control (QC) sample at 10x volume. Samples were vortexed for 10 seconds then snap frozen on dry ice. Thawed samples were sonicated for 15 minutes on ice, stored for 2 hours at -30^°^C, and then centrifuged for 15 minutes at 16,000×g (4^°^C). Supernatant was collected, aliquoted, dried down using a vacuum concentrator and stored at -80^°^C until LC/MS analysis.

Untargeted metabolomics was performed as published previously^18^ with slight modifications, using a 1290 Infinity II UHPLC coupled to a 6545 QTOF mass spectrometer via Dual AJS ESI source (Agilent, Santa Clara, USA) and MassHunter data acquisition software (v.10.1). Full scan MS data (m/z 50-1700) was acquired at a scan rate of 2.5 spectra/sec with the following source conditions: Gas temperature 250°C, gas flow 13 L/min, sheath gas temperature and flow at 400°C and 12 L/min, respectively, nebulizer 30 psi, fragmentor 135, capillary voltage at +4500 V and -4000 V, nozzle voltage was zero. Metabolite separation was performed on a Zorbax HILIC Plus RRHD (95Å, 1.8 µm, 2.1×100mm, Agilent) analytical column connected to a 3 × 5 mm Zorbax HILIC Plus UHPLC guard column. The autosampler and column temperature were set to 4°C and 40°C, respectively. In positive and negative mode, eluent A was 10 mM ammonium acetate in acetonitrile/milliQ water (95:5, v/v) and eluent B was 10 mM ammonium acetate in acetonitrile/milliQ water (50:50, v/v). The following gradient was used for both modes: 0 minutes (1% eluent B) - 3.5 minutes (50% B) - 5.5 minutes (99%B) - 6.5 minutes (99% B) - 6.7 minutes (1% B) - 12 minutes (1% B). Flow rate was set to 0.5 mL/min.

For LC/MS analysis, polar compounds were resuspended by addition of milliQ water, thoroughly vortexing and incubating for 10 min on ice. Following centrifugation, supernatant was transferred to a new tube and diluted with chilled acetonitrile containing 1 µg/mL Val-Tyr-Val internal standard to a final concentration of 80% acetonitrile. Injection volumes were 2 μL for positive and 3 μL for negative mode. Samples were run in a randomized order.

Positive and negative mode data was analysed separately using MassHunter Profinder (v 10 SP1, Agilent) recursive feature extraction method employing default settings with minor adjustments: Peak extraction was restricted to retention time (Rt) range 0-6.5 minutes, compound binning and alignment tolerances were set to 1% + 0.3 minutes for Rt and 20 ppm + 2 mDa for mass, integrator Agile 2 was used for peak integration, peak filters were set to at least 2500 counts and features must have satisfied filter conditions in at least 75 % of files in at least one sample group. Feature peak area was exported and data cleaning was performed using an in-house R script compiled of the following steps. Features were deleted if they: had a mean QC/tube blank area ratio of < 10; were absent across all QC samples; and had duplicates present. In addition, samples with a total ion current (TIC) scaling factor more than 50% above or below the median TIC were removed. Differential expression analysis and principal components analysis (PCA) were conducted using MetaboAnalystR 2.0 ^19^, including additional data pre-processing steps: Features with >50% missing values were removed, missing data values were imputed using k nearest neighbour (kNN), sample normalisation was performed by reference sample (probabilistic quotient normalization) and a log transformation was applied.

### *In vitro Pb*A maturation assay

*Pb*A infected blood was added to culture medium (RPMI, 5 U mL^-1^ heparin sulphate) at 2.5% haematocrit, containing 10% vol mouse plasma. Samples were cultured in 5% O_2_, 5% CO_2_, 90% N_2_ at 37^°^C for 22 hours.

### Droplet-based scRNA-seq of pRBC

Peripheral blood (5-6 drops) was collected into 1ml cold RPMI medium containing 5 U/mL heparin by tail vein bleed. Diluted blood (50 μL) was further diluted in 2 mL cold 1% BSA/PBS and RBCs isolated away from leukocytes by cell sorting. RBC, containing pRBC, were loaded such that ∼5,000 pRBCs were loaded per channel onto a Chromium Controller (10X Genomics) for generation of gel-bead-in-emulsions. Sequencing libraries were prepared using Single Cell 3’ Reagent Kits V3.1 for main experiment or V2 for repeat experiment (10X Genomics) and sequenced on a NextSeq550 (Illumina) using paired-end 150-base pair reads.

### scRNA-seq data quality control and analysis

FASTQ files were processed using Cell Ranger version 2.1.0 (10X Genomics) for the reference dataset and version 3.0.2 for the study of impaired maturation with combined genomes of mouse (mm10 - Genome Reference Consortium) and *Pb*ANKA (PlasmoDB release 39; ^13^) as a reference. ^20^. For quality control, RBC were first classified as containing only parasite signal, mouse or both. Using Cell Ranger all genes of mouse origin were filtered out. Seurat v3.2.2 “SCTransform” function was used for normalisation, finding of variable features, and scaling of data ^21^. Uniform manifold approximation and projection (UMAP) for dimension reduction (UMAP; ^22^) coordinates were generated using Seurat’s “RunUMAP” function based on 10-15 principal components (PCs). Differential gene expression analysis was performed with Seurat’s “FindMarkers” function using Wilcoxon-rank sum test, with all parameters kept at default except min.pct = 0.25. Integrated dimensionality reduction with the malaria cell atlas (MCA) datasets was performed using single-cell variational inference (scVI) version number v0.7.1^23^, with all parameters kept at default except dispersion parameter set as ‘gene-batch’. Each dataset was identified as separate batch. Computed latent variables were used as input to generate UMAP using Seurat’s “RunUMAP” function. To infer parasite life-stage from scRNA-seq data, PlasmoDB was used to extract *Pb*A genes from bulk RNA-seq data ^13^, upregulated at least 2-fold for each asexual blood-stage compared to the highest average gene expression value in the remaining two life-stages, resulting in lists of 291 genes for to ring-, 581 genes for trophozoites, and 754 genes for schizont stages. Mean expression of all genes in these genes signature lists was calculated for each cell and visualised using ggplot2 package. The scRNA-seq datasets generated during this study will be deposited at ArrayExpress.

### Statistics

Statistical analyses were performed using Prism 7 version 7.0c (GraphPad Software, San Diego, USA) or R version 3.6.3, the latter was used only for statistical analysis, one-way ANOVA, of transcript numbers from scRNAseq data. Time course graphs depict either individual data points or mean ± standard deviation (SD). Time course experiments were analysed using either two-way ANOVA multiple comparisons test, multiple comparisons test with a factor for time-point and a factor for treatment group (either Tukey or Dunn–Šidák corrected), or mixed-effect analysis as described in figure legend. Other analyses include one-way ANOVA and t-test as described in figure legend. A p value of 0.05 was considered significant. P values are shown as *p<0.05, **p<0.01, ***p<0.001, ****p<0.0001.

### Data availability

Mass spectrometry metabolomics raw and extracted feature data have been deposited to the NIH Common Fund’s National Metabolomics Data Repository (NMDR) website, the Metabolomics Workbench (https://www.metabolomicsworkbench.org) where it has been assigned Project ID PR0001195. The data can be accessed directly via it’s Project DOI (https://doi.org/10.21228/M87M55).

## Results

### Systemic host inflammation impairs maturation of blood-stage parasites *in vivo*

Previously, we used RBC adoptive transfer into mice to study maturation of a single cohort of blood-stage parasites^3-5,12,24^. Fluorescently-labelled RBC containing *Pb*ANKA-eGFP (defined as Generation 0 (Gen_0_)) are transferred into recipient mice, with flow cytometric enumeration and life-staging in peripheral blood every 4-6 hours. In this study, to determine if systemic host inflammation alone could impair parasite maturation *in vivo*, mice were pre-treated with Toll-like Receptor (TLR) 4, TLR9 or TLR3 agonists (LPS, CpG or Poly I:C) or control saline, prior to receiving a cohort of CellTrace™ FarRed (CTFR)-labelled RBC containing *Pb*ANKA-eGFP (Figure 1A). After parasites mature within Gen_0,_ they burst out of CFTR^+^ RBC (which constituted a minority of all RBC in the mice), and the majority of resulting merozoites infected CFTR^-^ RBC, defined as Generation 1(Gen_1+_)(Figure 1B). Therefore, this technique also permitted examination of parasite transit from one RBC to the next *in vivo*.

**Figure 1.**
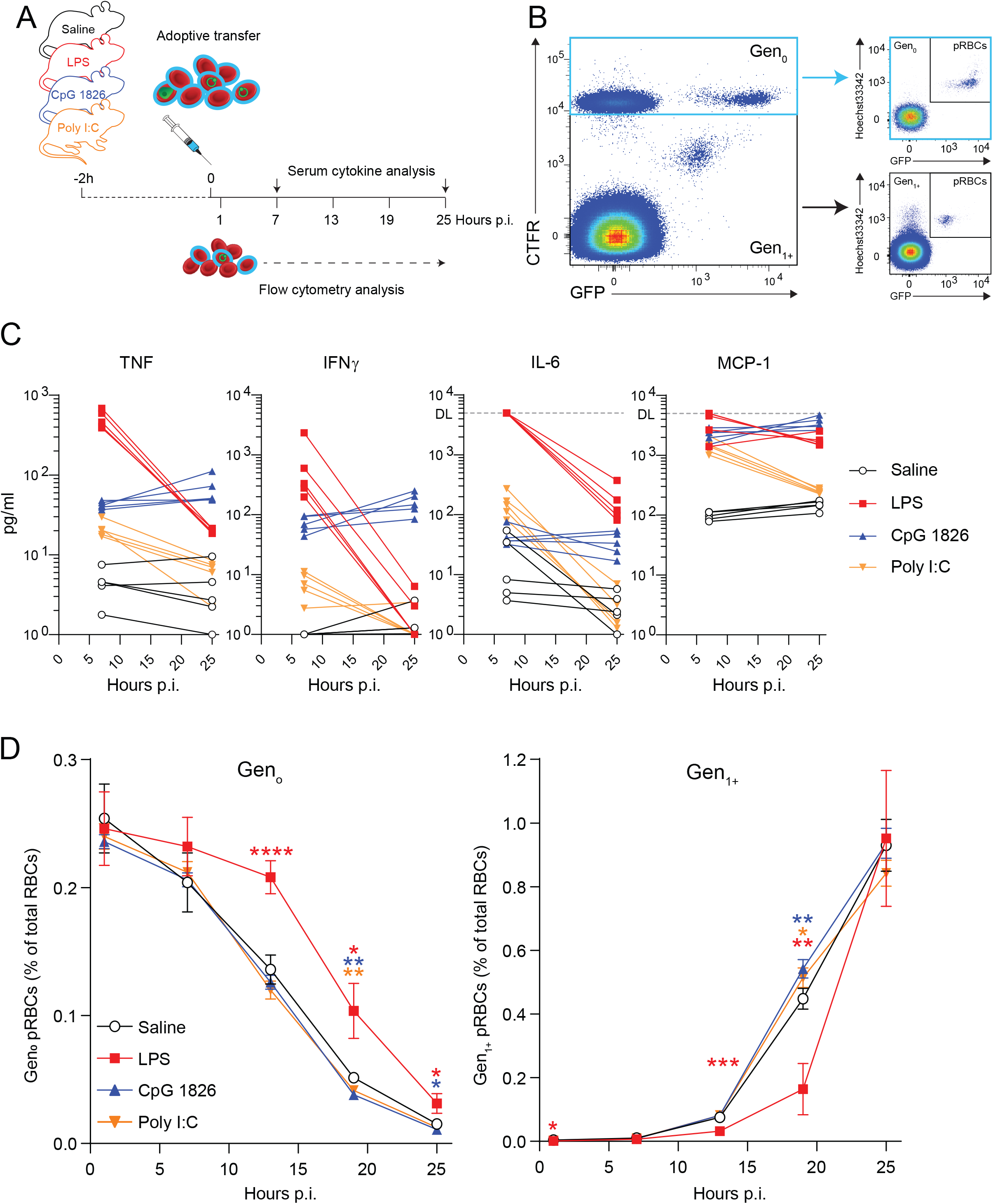
LPS-induced systemic inflammation impairs blood-stage parasites *in vivo*. A) Schematic of CTFR-labelled RBC (∼2-5% containing *Pb*A-GFP parasites) transferred into recipient mice (n=5/group), 2 hours after treatment with LPS (150 μg), CpG 1826 (50 μg) or Poly I:C (400 μg) or control saline, with peripheral blood assessed by flow cytometry at timepoints indicated. B) Representative flow cytometry gating for Gen_0_ (CTFR^+^) and Gen_1+_ (CTFR^-^) RBCs and assessment of parasites via Hoechst33342 and GFP expression. C) Serum cytokine levels tracked in individual mice at 7 and 25 hours post-infection - dotted line: assay detection limit (DL). D) Enumeration of Gen_0_ and Gen_1+_ pRBCs over time. Data presented as mean ± SD. Statistical significance tested relative to saline controls. Data representative of two independent experiments. Statistical analysis: two-way ANOVA with a factor for time-point and for treatment group. Testing for a treatment group effect in Gen_0_ (p<0.0001, F = 20.46, degrees of freedom (df) = 3) and Gen_1+_ (p=0.0087, F = 5.492, df = 3). *p<0.05, **p<0.01, ***p<0.001, ****p<0.0001 (Dunnett test for multiple comparisons).

At the doses employed all TLR agonists stimulated systemic cytokine production by 9 hours post-treatment, with significant increases in canonical inflammatory cytokines, TNF, IFNγ and Interleukin (IL)-6, and chemokine, MCP-1 (also known as CCL2)(Figure 1C). At the doses employed, differences were apparent between the agonists (Figure 1C). LPS triggered the largest systemic cytokine response at 9 hours post-treatment, CpG the most prolonged upregulation of TNF and IFNγ, while Poly I:C elicited the mildest response. We examined the fate of Gen_0_ and Gen_1+_ parasites by flow cytometry at 6 hourly intervals under these different conditions. As expected in saline-control mice ^12,24^, the majority of Gen_0_ parasites were gradually lost over 25 hours, through rupture and transition to Gen_1+_, with some host clearance also implicated ^3^. We observed no effect of CpG or Poly I:C treatment on the dynamics of Gen_0_ or Gen_1+_ parasites (Figure 1D). In contrast, LPS-treatment caused parasites to persist in Gen_0_, which was associated with a delay in the emergence of Gen_1+_ parasites (Figure 1D). These data revealed at the doses employed, that systemic LPS treatment, but not CpG or Poly I:C, modified the rate at which parasites transitioned from one RBC to the next *in vivo*. Given differences in magnitude of systemic cytokine responses elicited by each treatment, our data suggest that potent host inflammatory responses alone can modulate maturation rate of asexual blood-stage parasites.

### Systemic inflammation alters the metabolomic composition of host plasma

Since LPS-conditioning had caused parasites to persist in Gen_0_ (Figure 1), similar to that reported for wild-type mice with acute malaria-induced inflammation^12^, we next determined whether circulating Gen_0_ parasites had been exposed to altered host plasma environments in these two scenarios. We tested the hypothesis that LPS-conditioning and host inflammation during acute malaria had altered plasma metabolomes. We also examined 5-day infected and naïve *rag1*^*-/-*^ mice, since Gen_0_ parasite maturation is largely unimpeded in these hosts^12^. Untargeted metabolomics using liquid chromatography-coupled mass spectrometry (LC/MS) with either positive or negative ionisation was conducted on plasma samples from individual mice (Figure 2A). Abundance of metabolite features for each sample were analysed by Principal Component Analysis (PCA) and Uniform Manifold Approximation and Projection (UMAP), where proximity of samples can be interpreted as metabolomic similarity (Figure 2B). Firstly, plasma samples within the same experimental group clustered together (in both positive or negative mode), revealing as expected similar metabolomes amongst similarly-treated mice (Figure 2B). Consistent with our hypothesis, plasma metabolomes after 9 hours of LPS-conditioning, and in 5-day infected wild-type mice did not overlap with, and were located at a substantial distance from naïve wild-type controls (Figure 2B). Moreover, although plasma metabolomes were altered by acute malaria in *rag1*^*-/-*^ mice, an environment observed to induce little to no impaired maturation ^12^, this occurred to a lesser degree than in wild-type mice (Figure 2B). Results were replicated independently (Supp Figure 1). Thus, impaired maturation of Gen_0_ parasites, induced by systemic host inflammation, is associated with a substantial alteration in the host plasma metabolome.

**Figure 2.**
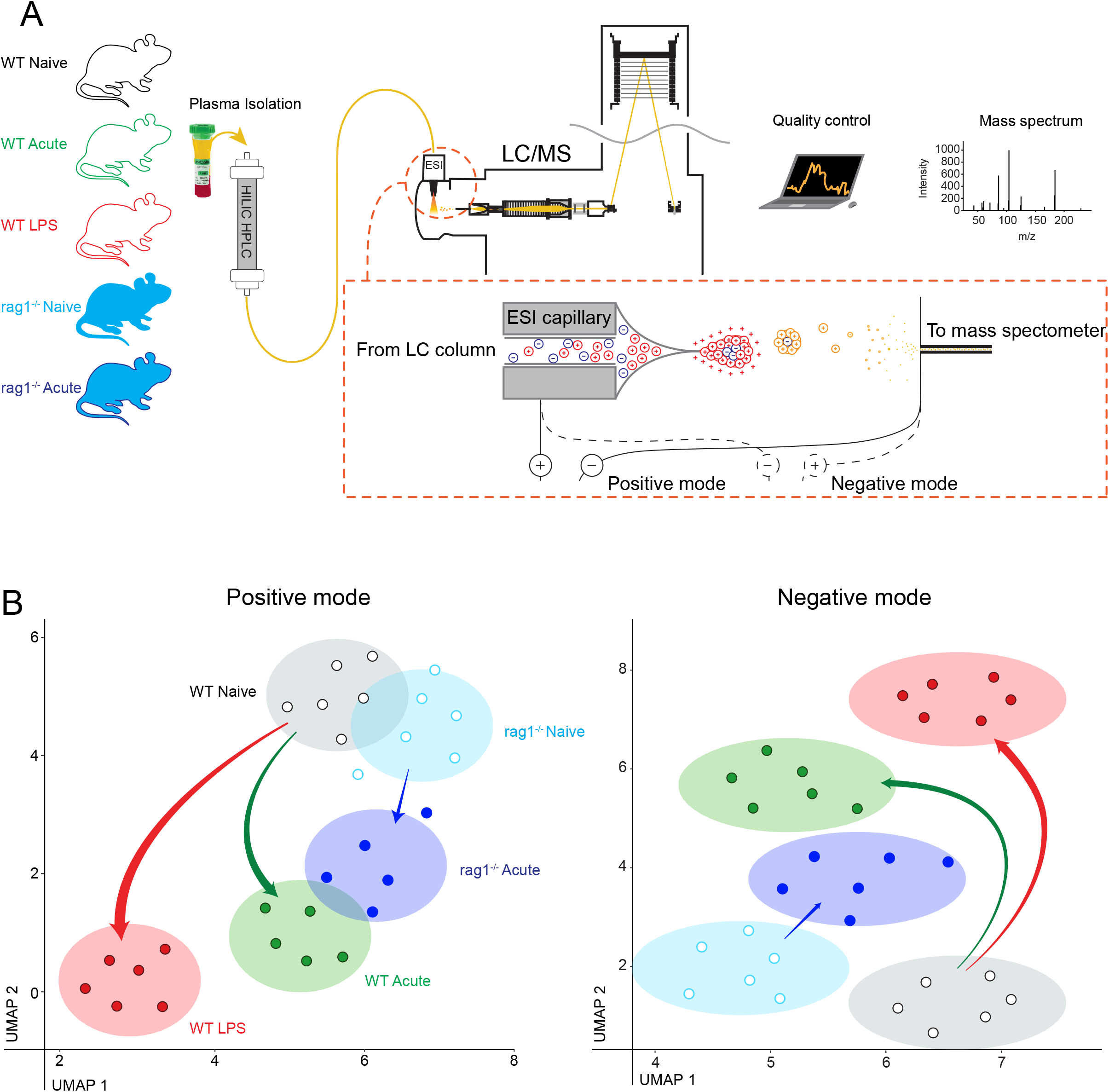
Systemic inflammation alters the metabolomic profiles of circulating plasma *in vivo*. A) Schematic showing plasma samples (n=6 per group) separated on a hydrophilic interaction liquid chromatography (HILIC) column, followed by ionisation and mass spectrometry. B) UMAP dimensionality reduction representation of untargeted LC/MS data obtained using positive or negative electrospray ionisation (ESI). Dots represent plasma metabolomes from individual mice, shaded ellipses depict centroid and 95% confidence intervals for each group. Arrow width indicates Euclidean distance between centroids of groups. Independent experimental repeat shown in Supplementary Figure 1.

### Host plasma conditioned by systemic inflammation directly impairs parasite maturation

To test for a causal link between changes in host plasma and impaired parasite maturation, we cultured parasites in medium supplemented with host plasma. We employed a previously reported *in vitro* maturation assay ^10^, focussing on the developmental progression that *P. berghei* ANKA parasites can complete *in vitro*. Flow cytometric assessment revealed that although ring/early trophozoites matured effectively in the presence of plasma from naïve mice, this was significantly impaired in the presence of plasma from mice either acutely infected or LPS-conditioned (Figure 3C). Moreover, the apparent defect in maturation could not be rescued by supplementation with naïve plasma (Figure 3D). It was also noted that schizont development, albeit modest, proceeded equally in all conditions (Figure 3E). Together, these data support a causal link between alterations to host plasma and impaired parasite maturation, specifically of ring and early trophozoite stages, but not for mature trophozoites progressing to schizonts. Moreover, our data implicate an unidentified inhibitory factor or factors, not nutrient starvation, as the mechanism underlying inflammation-induced impairment of ring/trophozoite maturation.

**Figure 3.**
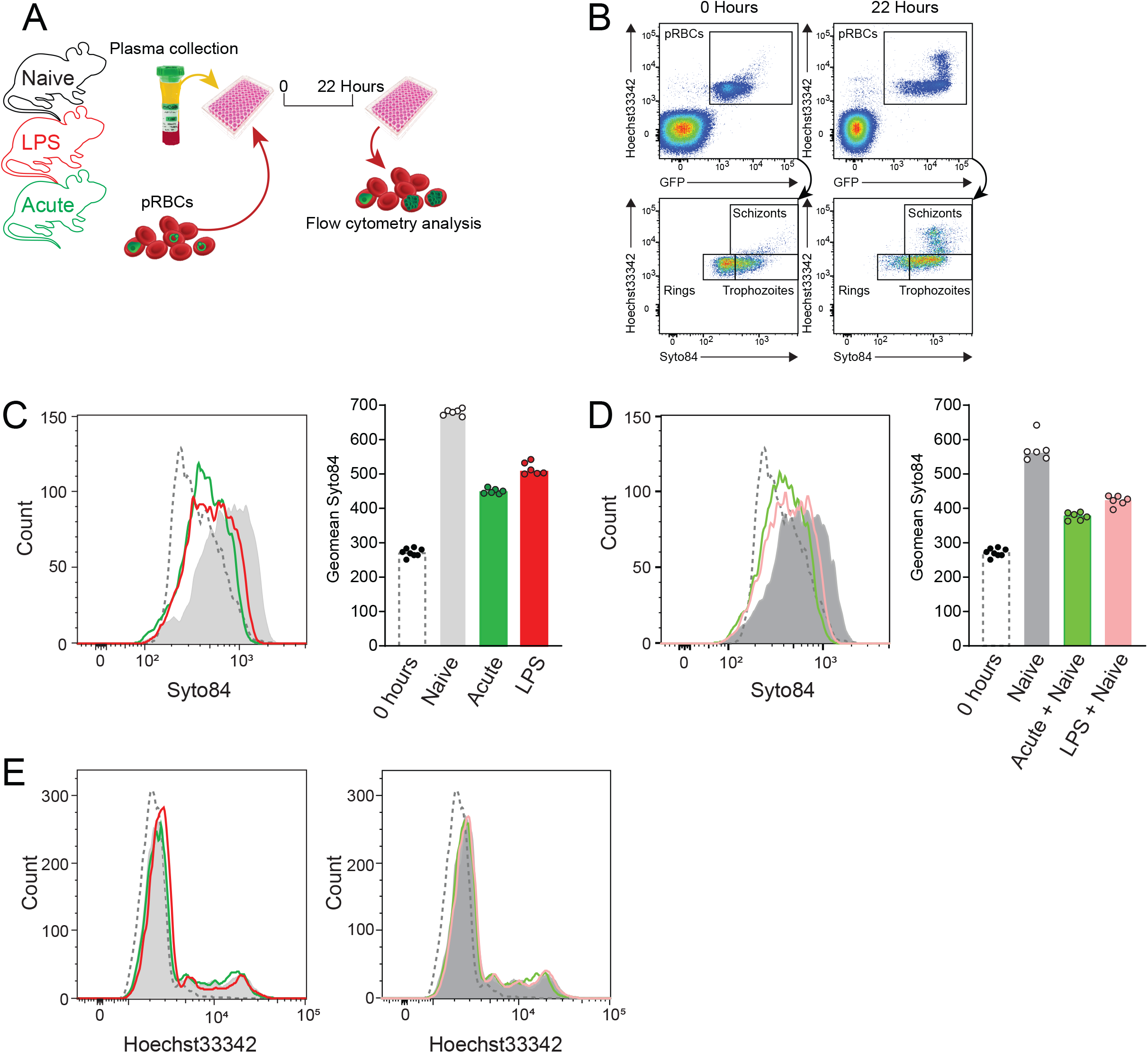
Plasma altered by systemic inflammation directly inhibits parasite maturation. A) Schematic showing pRBC from a passage mouse cultured *in vitro* for 22 hours, in media supplemented with pooled plasma from LPS-conditioned, acutely-infected or control mice (n=6/group), followed by flow cytometric assessment of maturation. B) Flow cytometry gating to identify pRBCs using Hoechst33342 staining and GFP expression, and life-stages using Hoechst33342/Syto84 staining at 22 hours p.i. compared to 0 hours. C) Representative histogram of Syto84 expression in non-schizont parasites after culture - 10% v/v plasma. Bar graph shows geometric mean Syto84 staining of technical replicates, data representative of two independent experiments. D) Representative histogram of Syto84 expression in non-schizont parasites after culture with plasma from LPS-conditioned or acutely-infected mice supplemented with equal volumes naïve plasma – 10% + 10% v/v plasma in all groups. Bar graph shows geometric mean of Syto84 staining in six technical replicates, data representative of two independent experiments. E) Representative histogram of Hoechst33342 staining in all pRBC prior to and after culture as in C) and D), with data representative of six technical replicate in two independent experiments.

### Droplet-based scRNA-seq maps asexual life-cycle progression of *P. berghei* ANKA parasites *in vivo*

To determine if parasites responded directly to altered plasma environments, we analysed parasites *ex vivo* via RNA-seq. Asynchronicity of *P. berghei* ANKA parasites *in vivo*, and possible differential effects on ring, trophozoite and schizont stages (Figure 3), necessitated a single-cell RNA-seq approach. Previous scRNA-seq work, presented in the Malaria Cell Atlas (MCA) ^15^, largely examined *P. berghei* after *in vitro* culture, where host inflammatory effects and host RNA particularly from recently invaded reticulocytes, would be absent.

To determine the feasibility of analysing individual parasite transcriptomes directly *ex vivo*, and to compare these data to the MCA, we first studied a *P. berghei* ANKA-infected *rag1*^*-/-*^ mouse (Figure 4A), for easier detection of all asexual life-stages including mature trophozoites and schizonts ^25^, and because host inflammatory effects would be reduced. After flow cytometric exclusion of white blood cells from whole blood, total RBC, including those parasitised, were processed for droplet-based 3’ scRNA-seq (Figure 4A). Dual mapping of scRNA-seq data to the *Plasmodium* and mouse genomes revealed the majority (79.8%) of RBC with detectable mRNA solely contained parasite-derived transcripts, while 3.7% contained host RNA alone (almost exclusively hemoglobin-encoding mRNA), or a mixture of hemoglobin and parasite mRNA (Figure 4B). These data suggested that direct *ex vivo* examination of parasites was not impeded by the presence of host RNA. We also noted similar numbers of parasite genes detected per RBC compared to the MCA (Figure 4C), confirming the sensitivity of our approach against this standard. To compare the quality of our direct *ex vivo* data to the MCA, which is itself was generated by two different methods (SMART-seq2 and 10x), we performed neural network-based integration of these three datasets using single-cell Variational Inference (scVI)^23^, as described before^26,27^, which highlights commonalities between datasets, removing technical batch effects. scVI-based integration revealed, as expected, no cross over between our *ex vivo* asexual stage data and the multiple sexual, liver and mosquito-stage parasites (Figure 4D). Instead, our data mapped closely with the asexual life-cycle stages of the MCA, and given the proportion of ring, trophozoite, and schizont stages present in our sample by flow cytometric analysis (Figure 4A), we noted the majority of direct *ex vivo* transcriptomes mapped onto MCA zones associated with ring and trophozoite stages, with a minority occupying the more mature schizont stages (Figure 4D). These data confirmed the feasibility of examining individual blood-stage parasite transcriptomes directly *ex vivo*, facilitating the screening of heterogeneous parasite populations after exposure to altered host plasma.

**Figure 4.**
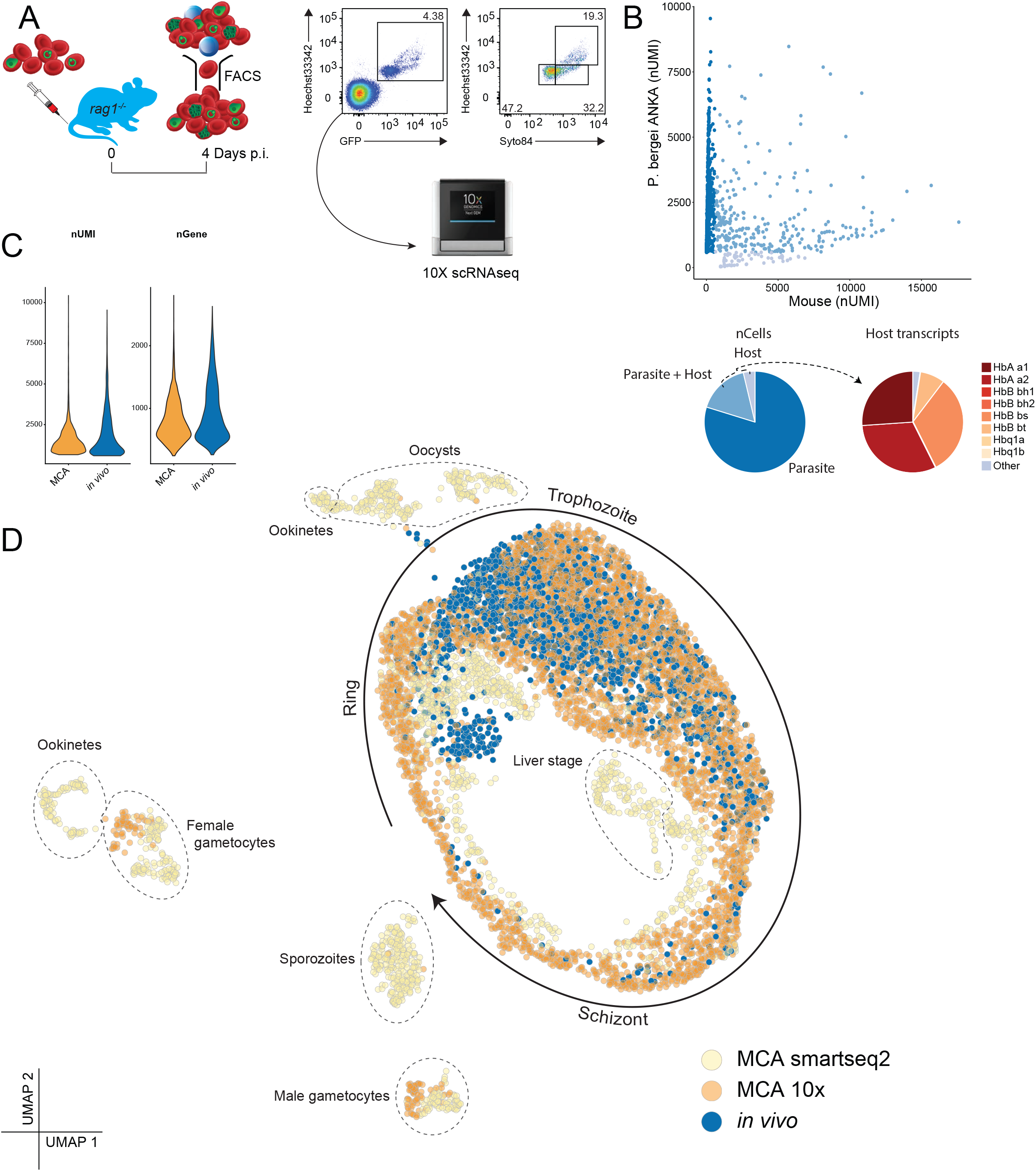
Droplet-based scRNA-seq of parasites directly *ex vivo*. A) Schematic for assessing droplet-based scRNA-seq of blood-stage parasites directly *ex vivo*. Blood from *Pb*A-GFP infected *rag1*^*-/-*^ mice were pooled (∼4.4% parasitemia, spectrum of life-stages as shown in FACS plots) prior to FACS sorting of RBCs and exclusion of leukocytes via FSC/SSC profiles. RBCs were loaded onto a Chromium Controller for transcriptomic analysis. B) Number of Unique Molecular Identifier (nUMI) transcripts mapping to mouse and parasite genomes for each cell. Pie charts show proportion of RBC classified as containing transcripts from parasites only, host only, or a mixture of the two (left), and for host transcripts, the proportions encoding haemoglobin genes (right). C) nUMI and number of genes detected per cell compared to Malaria Cell Atlas (MCA) blood-stage and gametocyte transcriptome. D) UMAP representation of scVI-integrated data of direct *ex vivo* blood-stages from this study with MCA 10x and SMART-seq2 data.

### Systemic host inflammation downregulates transcriptional and translational activity in trophozoites

To assess parasite transcriptomes directly *ex vivo*, we first reduced the time needed to FACS-sort transferred CFTR^+^ RBC from peripheral blood of LPS-conditioned, acutely-infected or naïve hosts (Figure 5A). Our published RBC adoptive transfer method^5,12,24^ results in ∼2-3% of all RBC in recipients being CFTR^+^ (Supp Figure 2A), a frequency incompatible with timely FACS-sorting within ∼20 minutes. So, we first confirmed that increasing the number of CFTR^+^ RBC injected per mouse by ∼5-fold elicited the same impaired maturation phenotype for Gen_0_ parasites as previously observed in acutely-infected mice (Supp Figure 2B).

**Figure 5.**
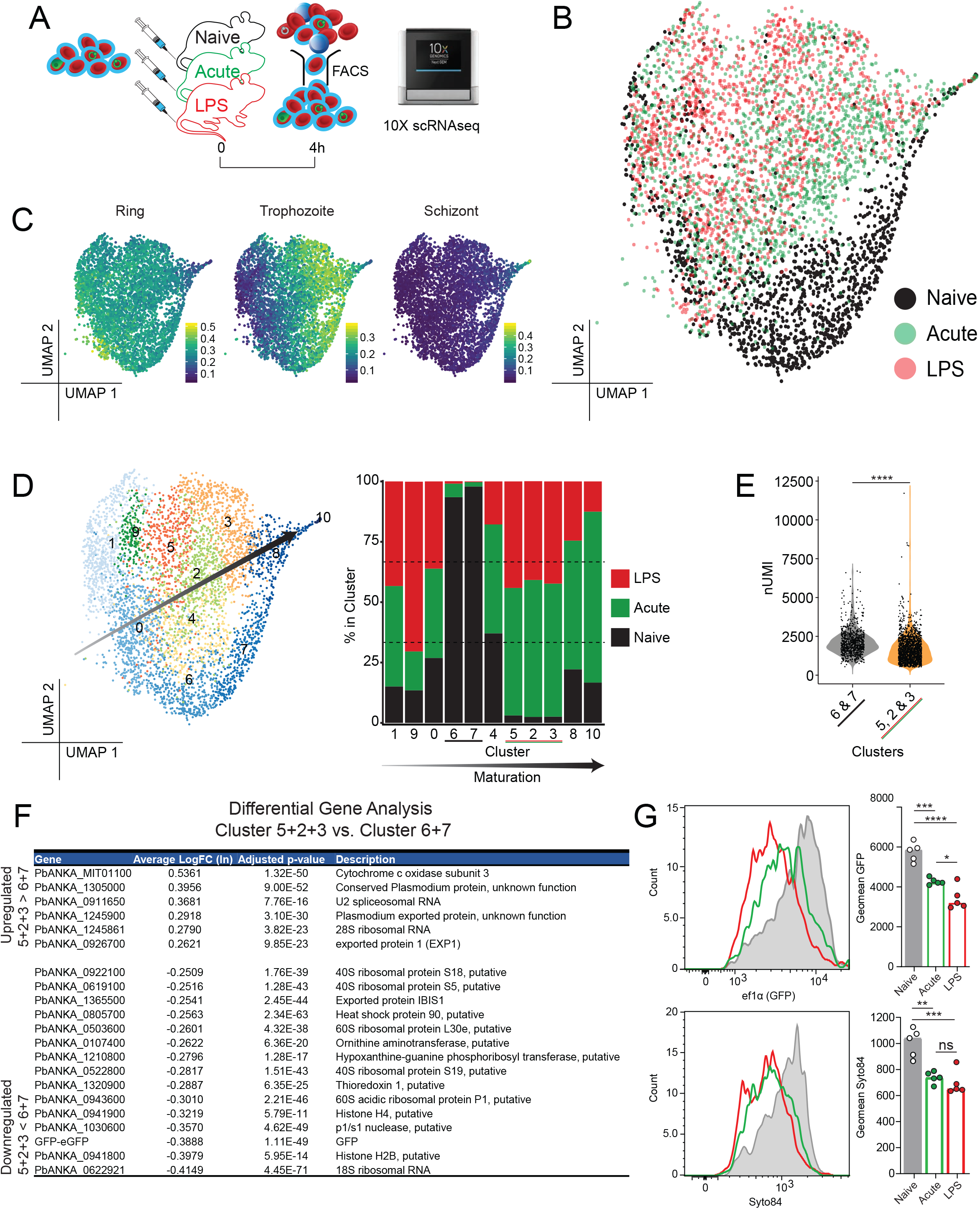
Rapid transcriptomic change in trophozoites exposed to systemic host inflammation *in vivo*. A) Schematic showing CFTR^+^ RBC (containing 4.8 % *Pb*A-GFP parasites) transferred into LPS-conditioned, acutely-infected or control mice (n=5/group), recovered by cell sorting after 4 hours, and immediately loaded onto a Chromium controller for scRNA-seq analysis. B) 2D UMAP after PCA of post-QC individual parasite transcriptomes after 4 hours *in vivo* exposure to host-inflammation. C) Expression of ring-, trophozoite- and schizont-gene signatures across the UMAP embedding taken from PlasmoDB (https://plasmodb.org/plasmo/app). D) Relative contribution of parasites from each group within each of 11 clusters, defined by unsupervised clustering on transcriptomic similarity; arrow indicates inferred rough directionality of maturation from ring to trophozoite to schizonts within the pooled data. E) Number of Unique Molecular Identifiers (nUMI) per cell for pooled clusters 6+7 and 5+2+3. F) Differentially-expressed genes between pooled clusters 5+2+3 versus pooled clusters 6+7 ranked by average Log_n_FoldChange. G) Representative histograms of GFP expression driven of the ef1alpha promoter (top) and Syto84 staining (bottom) in non-schizonts 13 hours post-transfer; bar graphs show geometric mean GFP and Syto84 in individual mice (n=5/group). Statistical analyses performed t-test for panel E and one-way ANOVA for G. E) Testing for a grouped cluster effect (p<0.0001, t=160.9, df=1). G) Testing for a treatment group effect in GFP (p<0.0001, F=34.65, df=2) and Syto84 (p=0.0002, F=18.35, df=2). *p<0.05, **p<0.01, ***p<0.001, ****p<0.0001. (Tukey test for multiple comparisons).

Next, we adoptively transferred this higher number of CFTR^+^ RBC (∼4% parasitised) equally amongst three different groups of conditioned hosts – LPS-conditioned, acutely-infected, or control naïve (Figure 5A). Given evidence of altered phenotypes in asexual parasites *in vivo* by 6 hours post-exposure ^10,12^, we recovered CFTR^+^ RBC after 4 hours, to capture early transcriptional changes in Gen_0_ parasites. In parallel, we also confirmed at subsequent time points, 13 and 25 hours post-transfer, that LPS-conditioning and acute infection had indeed caused Gen_0_ parasites to persist (Supp Figure 3A & B). Total CFTR^+^ RBC recovered at 4 hours from each of 5 mice per group were pooled in roughly equal proportions, and immediately loaded for droplet-based scRNA-seq, with the objective of capturing ∼3000 parasite transcriptomes per condition.

After mapping to the *P. berghei* ANKA genome, and removal of low-quality pRBCs and uninfected RBCs, we recovered a total of 4,772 parasites across all three conditions (1500 for LPS, 1970 for acute-infection, and 1302 for naïve). PCA followed by UMAP revealed a heterogeneous spectrum of parasite transcriptomes in naïve mice (Figure 5B). Strikingly, many transcriptomes from LPS-conditioned or acutely-infected mice appeared to occupy the same unique areas of the UMAP away from naïve transcriptomes (Figure 5B). Using publicly available gene signatures for ring, trophozoite and schizont stages from *PlasmoDB*, we inferred ring-stages dominated a minor portion on the left of the UMAP, with schizonts located towards an extreme right point of the data (Figure 5C). Consistent with our earlier assessment in *rag1*^*-/-*^ mice (Figure 4), and the MCA ^15^, ring and schizont transcriptomes were more uniform compared to trophozoite stages.

We inferred a spectrum of life-stages in our dataset, from youngest rings at the lower left of the UMAP, to mature schizonts at the upper right (Figure 5D). Importantly, host inflammation had diverted the transcriptomes of trophozoites, which we confirmed in an independent repeat scRNA-seq experiment for acute infection (Supp Figure 4 A-D; Cluster 0 versus 3). Unsupervised clustering based on transcriptomic similarity produced 11 clusters, with Clusters 0, 1 & 9 most ring-stage like, Clusters 8 & 10 most schizont like, and Clusters 2-7 trophozoite-like (Figure 5D). To quantify apparent stage-specific transcriptomic differences, we examined the proportion of transcriptomes in each Cluster from each of the three host conditions (Figure 5D). While ring and schizont-like clusters (0, 1, 8, 9 & 10) had representation from all three conditions, trophozoite Clusters 6 & 7 were almost entirely composed of transcriptomes from naïve mice, while trophozoite Clusters 2, 3 & 5 were dominated by transcriptomes from LPS-conditioning and acute infection (in equal measures). The number of unique mRNA molecules (nUMI) recovered per trophozoite was reduced in pooled Cluster (2+3+5) compared to Cluster (6+7) (Figure 5E), a phenomenon not observed for rings or schizonts (Supp Figure 3C). This suggested a quantitative reduction in mRNA content, and thus transcriptional activity, in trophozoites responding to systemic host inflammation. However, qualitative differences were also evident, with 15 genes significantly down-regulated by systemic host inflammation in trophozoites, and only 6 genes up-regulated (Figure 5F). The trend for systemic inflammation to down-regulate not up-regulate genes was also evident in the independent scRNA-seq repeat (Supp Figure 4E). Across both scRNA-seq experiments, the main recurring phenomenon was that 40% of down-regulated genes encoded ribosomal components, suggesting that systemic inflammation had reduced the protein translational capacity of trophozoites (Figure 5F and Supp Figure 4F). Consistent with scRNA-seq observation of down-regulation of the parasite transgene eGFP driven off the translation-associated elongation factor 1α promoter (Figure 5G and Supp Figure 4G), we noted by flow-cytometry at 13 hours post-transfer that translated eGFP protein was reduced in trophozoites by systemic host inflammation, and total RNA content, assessed by Syto84 staining, was also reduced (Figure 5G and Supp Figure 4G). Thus, our data reveal a short 4 hour period of *in vivo* exposure to systemic host inflammation reduces transcriptional activity and translational capacity in trophozoite-stage asexual parasites, which ultimately associated with reduced maturation and progression to the next generation of RBC.

## Discussion

Recent studies provide emerging evidence that the maturation rate of blood-stage *Plasmodium* parasites is not constant *in vivo*, but can be adjusted, for example, by nutrient starvation or altering circadian rhythms of the host ^10,11^. Similarly, we previously reported host-dependent impaired maturation of parasites ^12^. Here, we extend those findings with evidence that host inflammation altered the plasma environment surrounding parasites *in vivo*. After only 4 hours of circulation in an inflamed host environment, the parasites’ adjusted their transcriptomic profile and protein translational capacity specifically in trophozoites, resulting overall in a lower rate of maturation within RBC. To the best of our knowledge, this is the first study to demonstrate that the transcriptomes of individual parasites can be modulated by inflammation within a host. This result further emphasises the dynamic nature of host-parasite interactions in this infection model.

We demonstrated with LPS-conditioning, that systemic host inflammation alone, in the absence of confounding factors such as ongoing infection ^12^, slowed the rate at which parasites transited from one generation of RBC to the next. While this is consistent with the idea that host inflammatory responses can impair parasite maturation, it was intriguing that other TLR agonists, CpG and Poly I:C, did not elicit such a response. Based on the range of serum cytokine responses elicited, the magnitude of inflammatory response may be key in delaying maturation of parasites. However, whether proinflammatory cytokines themselves might have direct anti-parasitic effect remains unresolved, although at least for TNF appears unlikely^28^.

Untargeted metabolomics permitted an unbiased assessment of the small molecule composition of plasma in which parasites circulated. We observed a clear perturbation of the plasma metabolome by either LPS-conditioning or acute malaria, both of which elicited impaired maturation *in vivo*. Furthermore, *in vitro* culture with plasma from LPS-conditioned or acutely infected wild-type mice directly impaired maturation of parasites prior to progression to schizont stages, which, given the failure of normal plasma to restore normal maturation, was consistent with inhibitory factors being present in altered plasma. UMAP analysis indicated that LPS-conditioning elicited a qualitatively distinct plasma metabolome compared to acute infection. Hence, it is possible that each condition mediated impaired maturation via distinct pathways, despite triggering similar changes to trophozoite transcriptomes. The precise identity of inhibitory factors induced by host inflammation remain to be identified. Previous studies have suggested that elevated lactate levels may be inhibitory for malaria parasites ^6,7^. However, we were unable to replicate these observations *in vitro* (data not shown). Future studies, using LC-MS/MS will be required to define the inhibitory factors in plasma induced by host inflammation. We expect that comparison of multiple scenarios in which impaired maturation either does or does not occur will facilitate this identification process.

In this study we hypothesized that metabolomic changes to the host plasma during systemic inflammation could be sensed and responded to by the parasite itself. Transcriptomic methods such as mRNA-sequencing (RNA-seq) offer a systems-wide view of unicellular and multi-cellular eukaryotic organisms. In particular, cellular change over a period of hours is captured well using RNA-seq. In our experimental system, *P. berghei* is asynchronous, meaning that all stages of the blood-stage life cycle are present at any given time. The heterogeneity of parasites, and the possibility of host mRNA contamination *in vivo* necessitated the adaptation of recent *Plasmodium* scRNA-seq techniques ^15^. In particular, it was necessary to remove peripheral blood mononuclear cells from our blood samples, achieved by flow cytometric sorting prior to droplet-based scRNA-seq. In addition, while host mRNA was detected in some RBCs, these encoded for hemoglobin almost exclusively, and constituted only a small minority in our data. By successfully integrating our *in vivo* data with *in vitro* generated scRNAseq data held within the MCA ^15^, we concluded that *in vivo* scRNA-seq analysis of *Plasmodium* parasites was feasible. Interestingly, we noted that as in the MCA, that while rings are transcriptomically highly homogeneous, and likewise schizonts, trophozoites display comparatively higher levels of heterogeneity. It is intriguing to consider what factors might control the differentiation trajectory of parasites as the progress from ring to schizont stages. Indeed, whether this transcriptomic variation is functionally relevant remains unclear, although it perhaps indicates that trophozoites more so than other forms, are adaptable. We expect future studies will benefit from this capacity to study individual parasites in fine detail. Importantly, by using this technique, we observed a specific effect of host inflammation, not on rings or schizonts, but on trophozoites. Moreover, we noted specific effects on genes related to protein translation, suggesting a molecular mechanism underpinning delayed maturation of parasites. However, it was noted that the number of genes differentially expressed as a result of host inflammation was relatively small, although a genome-wide bulk effect on transcription was apparent (validated by flow cytometry). An attempt to compare the early effect of host-inflammation versus antimalarial drug treatment (with rapid acting, sodium artesunate) on parasite transcriptomes was not fruitful (data not shown), since four hours of *in vivo* artesunate treatment prevented mRNA recovery from parasites, perhaps suggestive of rapid killing. Nevertheless, this is consistent with host inflammation inducing maturation delay, but not killing of parasites. In addition, while our study was designed to detect early changes, assessments at later timepoints might reveal other transcriptional pathways modulated by host inflammation.

It should be recognised that although scRNA-seq can detect cellular change over moderate time frames of hours, rapid or even instantaneous cellular response can be undetectable. For example, differential phosphorylation of proteins, protein trafficking, or influx of calcium ions may be biologically crucial, and yet could be invisible to scRNA-seq. Future experiments could, for example, explore the phosphoproteome of parasites to provide a fuller picture of how parasites sense and respond to host inflammation.

Finally, we consider broader implications of our findings. Firstly, we propose that host inflammation can reduce parasite population growth via an under-appreciated mechanism, that of reducing parasite maturation rate, not active clearance or killing of parasites in the spleen. The phenomenon of developmentally arrested parasites is not unprecedented, however, since certain *Plasmodium* species have been reported to enter dormancy^29^, either as part of their natural life-cycle, or as a result of antimalarial drug treatment. Nevertheless, metabolite-mediated inhibition of parasites may constitute a new method for combatting parasite growth *in vivo*. Another interpretation of the phenomena presented here is that parasites actively sense an inhospitable host environment, and as a result trigger a stress-like coping mechanism. We speculate that inflammation-driven alterations to parasite maturation could also contribute to variation between parasites in symptomatic versus asymptomatic infections in endemic regions^30^. Our *in vivo* data, combined with previous reports support a model in which blood-stage *Plasmodium* parasites are acutely aware of their surroundings in the bloodstream, and can respond to a highly dynamic host environment. This likely imbues *Plasmodium* parasites with the means to remain adaptable and viable within a mammalian host.

## Supplementary Figure Legends

**Supplementary Figure 1.**
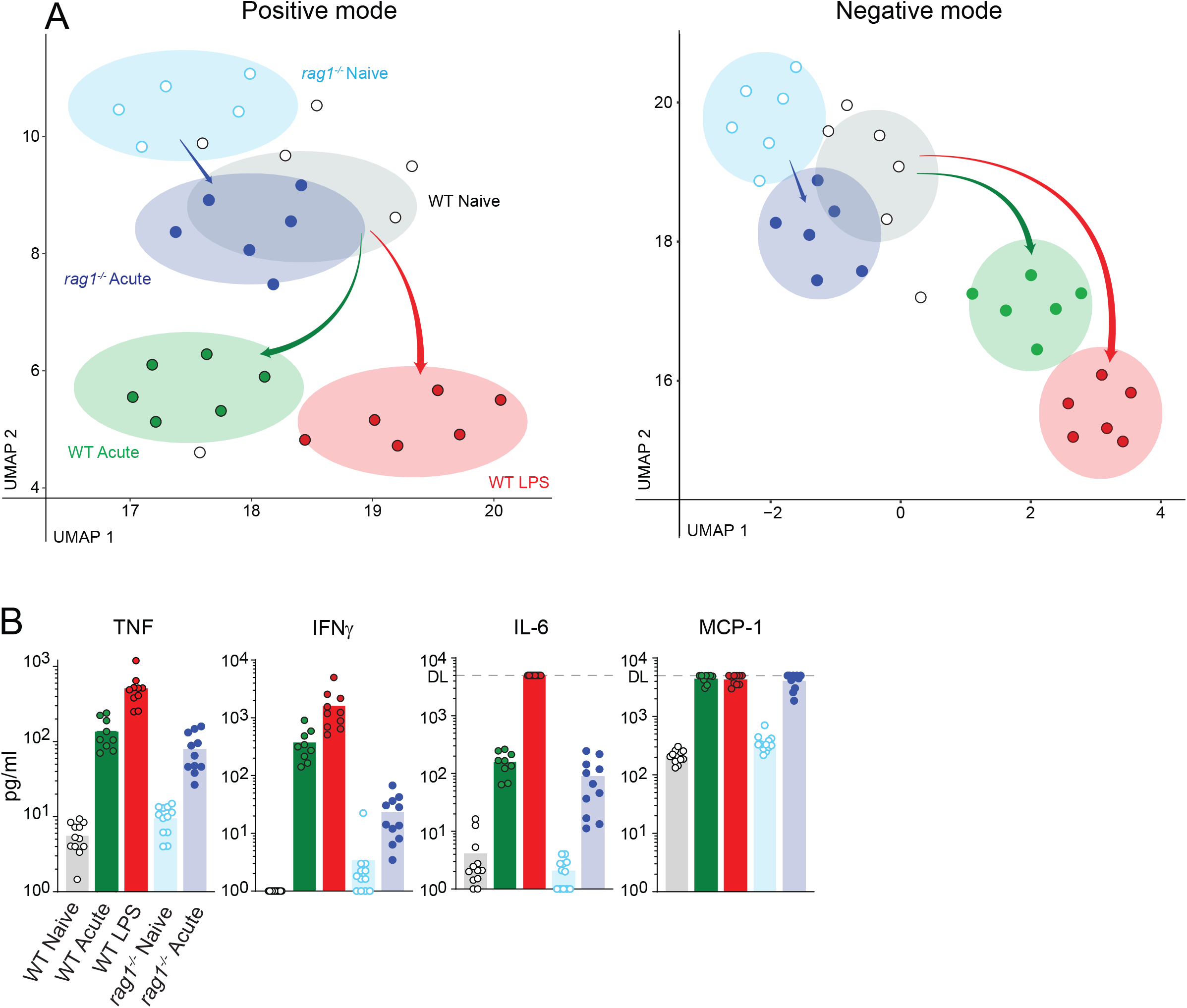
Experimental repeat of plasma metabolomic and cytokine assessments after LPS conditioning or during acute infection. A) UMAP dimensionality reduction representation of untargeted LC/MS data obtained for an independent repeat of data in Figure 2, using positive or negative electrospray ionisation (ESI). Dots represent plasma metabolomes from individual mice, shaded ellipses depict centroid and 95% confidence levels for each condition; arrow width indicates Euclidean distance between centroids of groups. B) Plasma cytokine levels in individual mice at time of metabolomic assessment, pooled from two independent repeat experiments (n=5 mice per group per experiment); bars represent mean value (n = 6 mice per group per condition per experiment); dotted line represents the assay detection limit (DL).

**Supplementary Figure 2.**
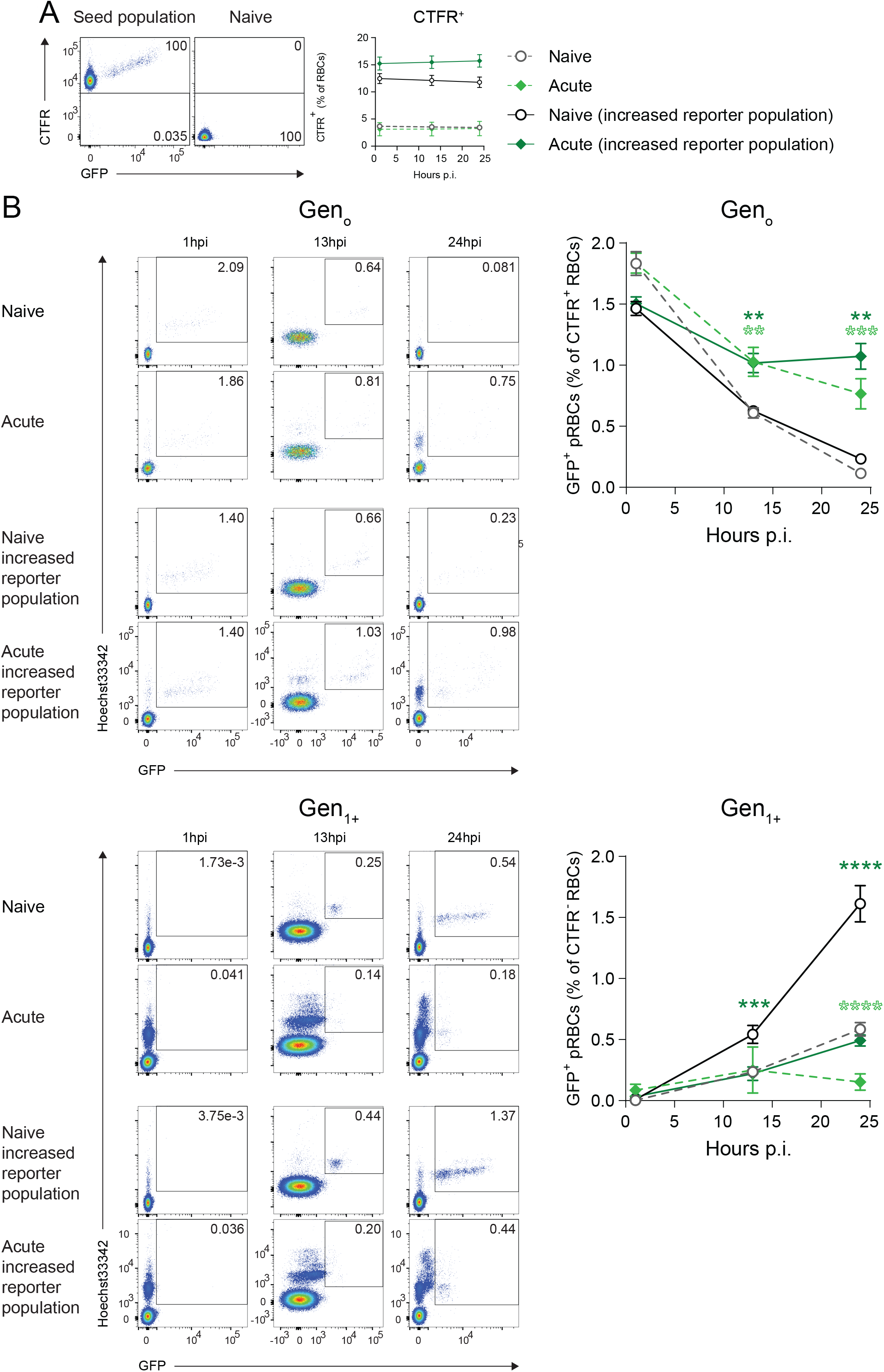
Increasing by 4-5-fold the number of CFTR^+^ RBC transferred into mice does not alter the impaired maturation phenotype. A) Flow cytometry plots showing CFTR^+^ RBC compared to naïve RBC; graph shows CFTR^+^ RBC dynamics as % of total RBCs for naïve or acutely-infected mice (n=5/group) receiving either the standard number or a 4-5 fold increased number of CFTR^+^ RBC. B) Representative flow cytometry plots of Gen_0_ and Gen_1+_ pRBCs expressed as a percentage of CFTR^+^ RBCt (n=5/group). Statistical significance indicated compared to appropriate naïve control. Statistical analysis: two-way ANOVA with a factor for time-point and for treatment group. Testing for a treatment group effect in Gen_0_ (p<0.0001, F=83.78, df=3) and Gen_1+_ (p<0.0001, F=113.5, df=3). *p<0.05, **p<0.01, ***p<0.001, ****p<0.0001. (Tukey test for multiple comparisons).

**Supplementary Figure 3.**
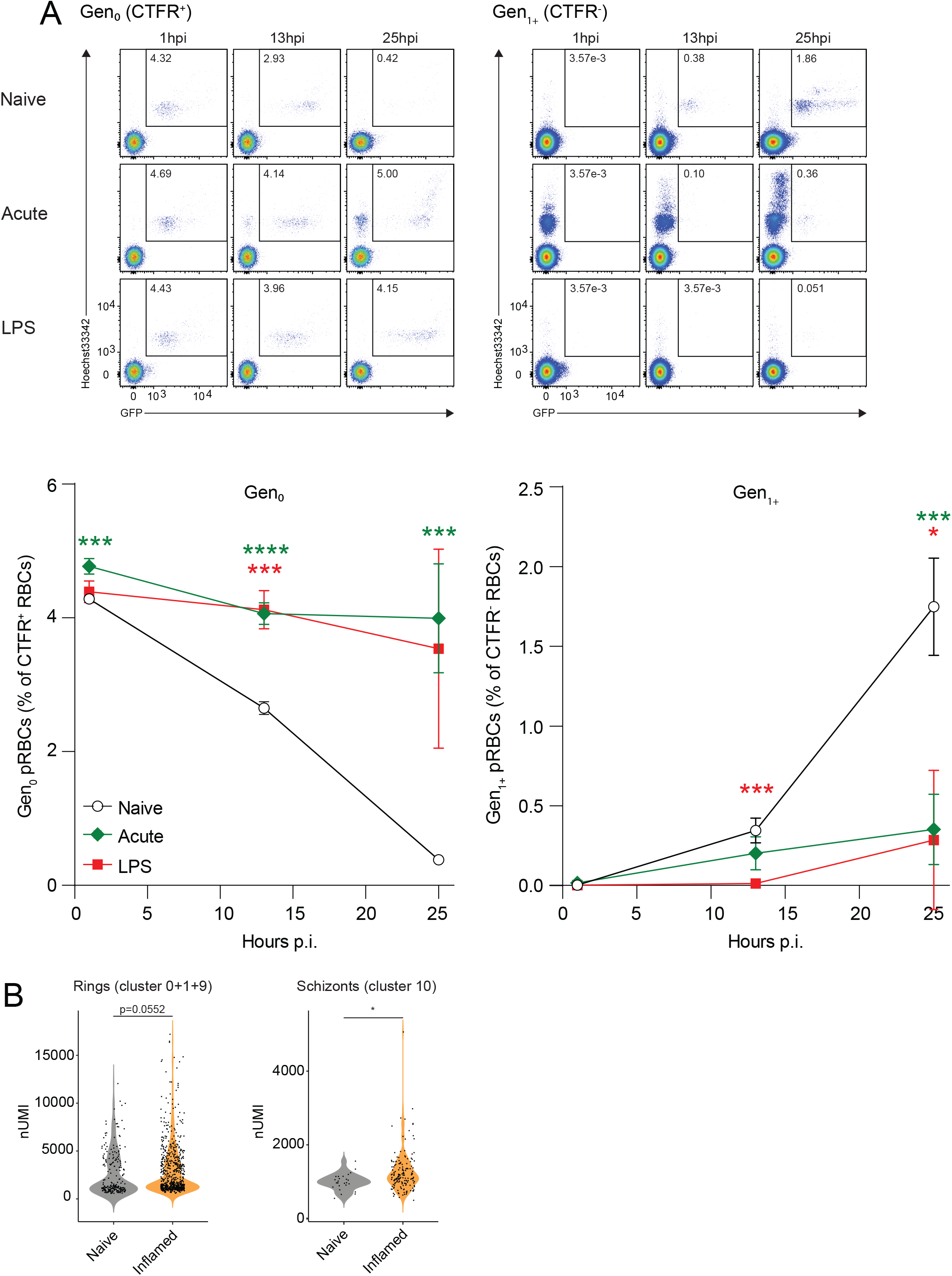
Confirmation of impaired maturation during the scRNA-seq experiment. A) Flow cytometry plots and B) summary graphs of Gen_0_ and Gen_1+_ parasites over time in LPS-conditioned, acutely-infected and control mice (n=5/group). Statistical significance indicated as compared to saline control. C) parasite nUMI per RBC for ring-stages or schizonts in naïve versus pooled acutely-infected and LPS-conditioned groups. Statistical analysis performed mixed-effects analysis for panel B and t=test for C. B) Testing for a treatment group effect in Gen_0_ (p<0.0001, F=55.06) and Gen_1+_ (p<0.0001, F=41.43). C) Testing for a condition effect in rings (p=0.0552, t=3.687, df=1) and schizonts (p=0.0157, t=5.938, df=1). *p<0.05, **p<0.01, ***p<0.001, ****p<0.0001. (Dunnett’s test for multiple comparisons).

**Supplementary figure 4.**
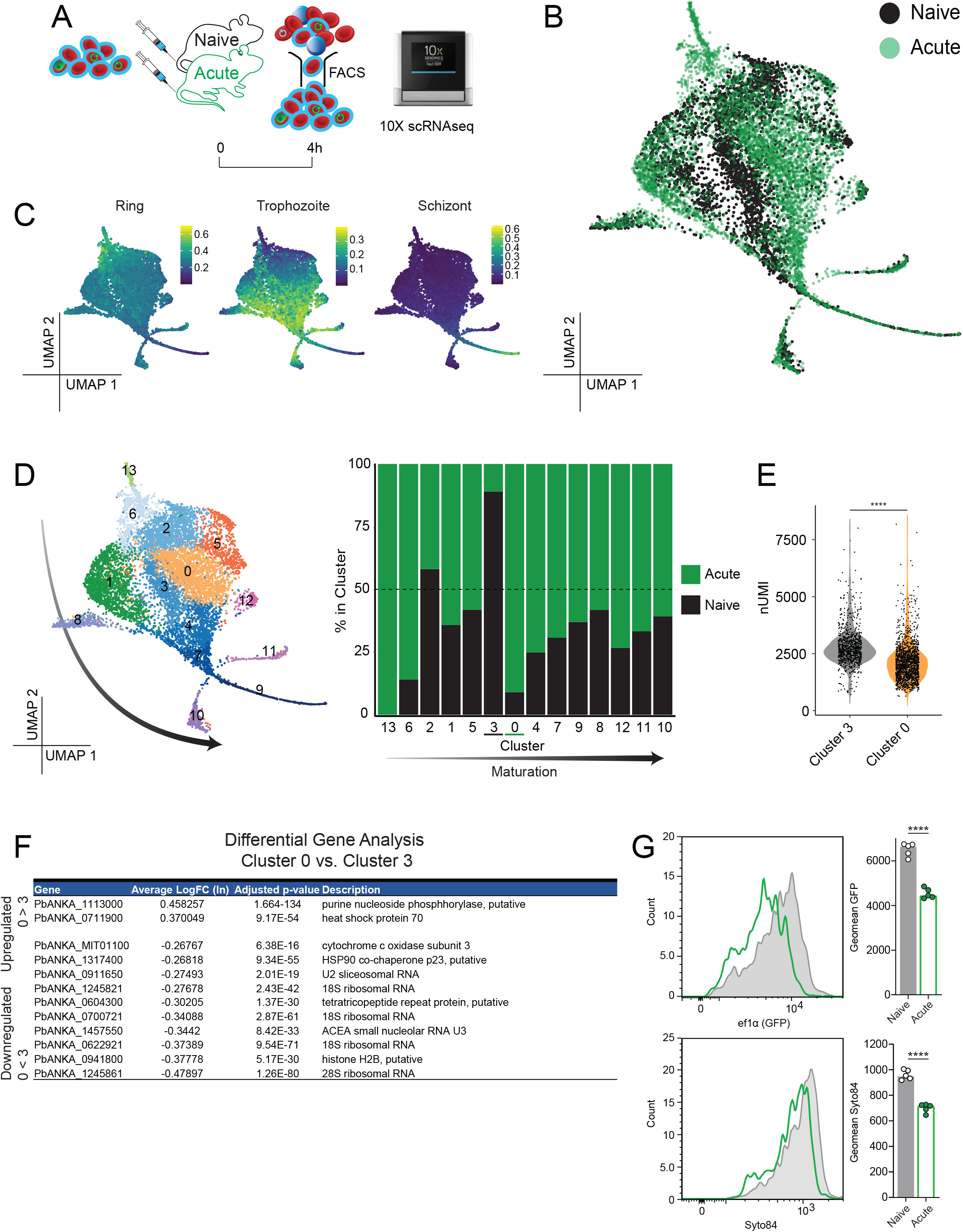
Experimental repeat of scRNA-seq assessment of parasites exposed to host inflammation. A) Schematic showing CFTR^+^ RBC (containing 3.7% *Pb*A-GFP parasites) transferred into acutely-infected or control mice (n=5/group), recovered by cell sorting after 4 hours, and immediately loaded onto a Chromium controller for scRNA-seq analysis. B) 2D-UMAP after PCA of parasite transcriptomes. C) Expression of ring-, trophozoite- and schizont-gene signatures across the UMAP embedding, taken from PlasmoDB (https://plasmodb.org/plasmo/app). D) Relative contribution of parasites from either group within each of 14 clusters, defined by unsupervised clustering on transcriptomic similarity; arrow indicates inferred rough directionality of maturation from ring to trophozoite to schizonts within the pooled data. E) Number of unique molecular identifiers (nUMI) per cell for cluster 3 and cluster 0. F) Differentially-expressed genes between cluster 0 versus 3 versus ranked by average Log_n_FoldChange. G) Representative histograms of GFP expression driven of the ef1alpha promoter (top) and Syto84 staining (bottom) in non-schizonts 13 hours post-transfer; bar graphs show geometric mean GFP and Syto84 in individual mice (n=5/group). Statistical analyses performed using t-test. E) Testing for a grouped cluster effect (p<0.0001, t=391.9, df=1). G) Testing for a treatment group effect in GFP (p<0.0001, t=11.78, df=8) and Syto84 (p<0.0001, t=11.38, df=8) *p<0.05, **p<0.01, ***p<0.001, ****p<0.0001.

